# Tissue disorder strength measured by quantitative phase imaging as intrinsic cancer marker

**DOI:** 10.1101/240457

**Authors:** Masanori Takabayashi, Hassaan Majeed, Andre Kajdacsy-Balla, Gabriel Popescu

**Affiliations:** Department of Systems Design and Informatics, Kyushu Institute of Technology, Iizuka, Fukuoka, Japan; Department of Electrical and Computer Engineering, Beckman Institute for Advanced Science and Technology, University of Illinois at Urbana-Champaign, Champaign, Illinois, United States of America; Department of Bioengineering, Beckman Institute for Advanced Science and Technology, University of Illinois at Urbana-Champaign, Champaign, Illinois, United States of America; Department of Pathology, University of Illinois at Chicago, Chicago, Illinois, United States of America

## Abstract

Tissue refractive index provides important information about morphology at the nanoscale. Since the malignant transformation involves both intra- and inter-cellular changes in the refractive index map, the tissue disorder measurement can be used to extract important diagnosis information. Quantitative phase imaging (QPI) provides a practical means of extracting this information as it maps the optical path-length difference (OPD) across a tissue sample with sub-wavelength sensitivity. In this work, we employ QPI to compare the tissue disorder strength between benign and malignant breast tissue histology samples. Our results show that disease progression is marked by a significant *increase* in the disorder strength. Since our imaging system can be added as an upgrading module to an existing microscope, we anticipate that it can be integrated easily in the pathology work flow.

## Introduction

Breast cancer is the most commonly diagnosed type of cancer among women worldwide [1]. Furthermore, according to the American Cancer Society, the incidence of breast cancer in the US is on the rise, with 200,000 new cases expected in the year 2017 [2]. While the burden of disease is significant, standard breast histopathology still relies on manual microscopic inspection of Hematoxylin and Eosin (H&E) stained tissue. The H&E primary stain provides the necessary contrast needed for a trained pathologist to distinguish between normal and abnormal tissue morphology. However, this type of investigation is qualitative, depends on the details of tissue processing and, as a result, often leads to inter-observer variability. Thus, there is a need to provide an objective basis for evaluation based on physical metrics. For cases where information provided by the H&E stain is limited and diagnosis is difficult, specialized stains can help pathologists [3]. New quantitative markers can provide objective assessment, as well as information complementary to traditional biomarkers. Specific information on tumor cell biology, extracted by such intrinsic markers, can also potentially lead to automated computer algorithms.

Quantitative phase imaging (QPI) is a label-free microscopy technique where contrast is generated by the optical path-length difference (OPD) across a tissue specimen [4–7]. The phase image (*x*, *y*) extracted in QPI is given by the expression

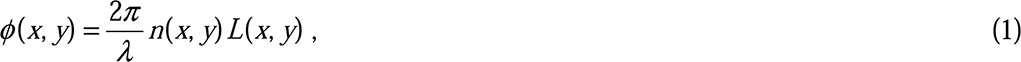
where *n*(*x*, *y*) is the refractive index contrast between the tissue and the surrounding medium (mounting medium in the case of histology samples), *L*(*x*, *y*) is the tissue thickness and *λ* the illumination wavelength [4]. For precise histological sections, the tissue thickness can be assumed to be relatively constant, *L*(*x*, *y*) ≈ *L*, meaning that a QPI measurement provides a signal that is proportional to the refractive index map of tissue. Since it is proportional to the dry mass content of cells and cellular matrix, the refractive index map informs on tissue density as well as cell organization within tissue [8, 9]. Since QPI allows extraction of the refractive index map label-free, the extracted biological markers are intrinsic, meaning that the results are not susceptible to variation due to differing staining procedures, thus, providing a robust signal for automated analysis. Tissue refractive index based markers have been used in the past to separate benign and malignant prostate tissue [10] as well as for detection of pre-malignancy in colorectal tissue [11]. OPD maps in general, extracted using QPI, have been used for addressing quantitative histopathology problems in prostate, colon, breast, pancreatic and other cancers etc. [12–20].

The tissue metric referred to as “disorder strength” was first used for diagnosis by Subramanian *et al*. [21]. The authors used it as a means of probing the sub-wavelength spatial fluctuations of refractive index and, thus, to detect carcinogenesis undetectable by standard histopathology [22–31]. Since QPI systems employ interferometric measurements, they are sensitive to sub-wavelength fluctuations in the refractive index map in both space and time [4]. Eldridge *et al*. measured the disorder strength using QPI and demonstrated that a transformation in cell mechanical properties can be measured by quantifying the cell disorder strength [32]. They applied this analysis to colon, skin and lung cancer cells to demonstrate an inverse relationship between sheer stiffness and disorder strength [32].

Here, we propose to use the disorder strength as an intrinsic marker for classifying benign and malignant breast tissue. We imaged a tissue microarray (TMA) comprising of cores obtained from cancer and normal-control patients. Details of this sample can be found in [13]. Each core was diagnosed as either benign or malignant by a board certified pathologist by examining H&E stained tissue images of a parallel tissue section. From this TMA, we studied 20 benign cores and 20 malignant cores. All research protocols involving the specimens were carried out with the approval of the University of Illinois at Urbana Champaign (UIUC) Institute Review Board (IRB Protocol Number 13900).

Since malignancy causes changes in both tissue architecture at the nanoscale, we hypothesize that these modifications will be reflected in the disorder strength. We demonstrate this by imaging the tissue microarray using called Spatial Light Interference Microscopy (SLIM), which is a high-sensitivity QPI method [5]. We detect sub-nanometer refractive index fluctuations and classify the tissue with high accuracy.

## Methods

A schematic of the SLIM setup is shown in Fig. 1A. The SLIM module is attached to a commercial phase contrast microscope (PCM). The lamp filament is imaged onto the condenser annulus (Köhler illumination conditions) which is located at the front focal plane of the condenser lens. The specimen is located at the back focal plane of the condenser lens, and front focal plane of the objective. The scattered and unscattered lights are relayed by the objective and tube lenses. As a result, the expanded phase contrast image which has the intensity distribution in accordance with the phase contrast caused by the specimen is observed at the image plane. However, because the output of PCM is the qualitative phase image, the quantitative phase map caused by the specimen cannot be directly retrieved from this image. The function of SLIM module is to convert this qualitative phase image into a quantitative one by properly phase modulating the incident light with respect to the scattered light. The field at the image plane is Fourier transformed by the lens L1, such that the unscattered light can be spatially isolated from the scattered light. Since the unscattered light has the ring form, by displaying the corresponding ring pattern on the reflective liquid crystal phase modulator (LCPM), we insure that the scattered light remains unaffected. Four phase shifts are applied to the unscattered light at increments of π/2 rad. as shown in Fig. 1B. The corresponding four images captured by the charge coupled device (CCD) are obtained as shown Fig. 1C. Consequently, the quantitative phase image is retrieved as described in Ref. [5] and Fig. 1D. Figure 2A and 2B shows the quantitative phase image and its expanded view of benign and malignant breast tissue samples, respectively.

**Fig 1.**
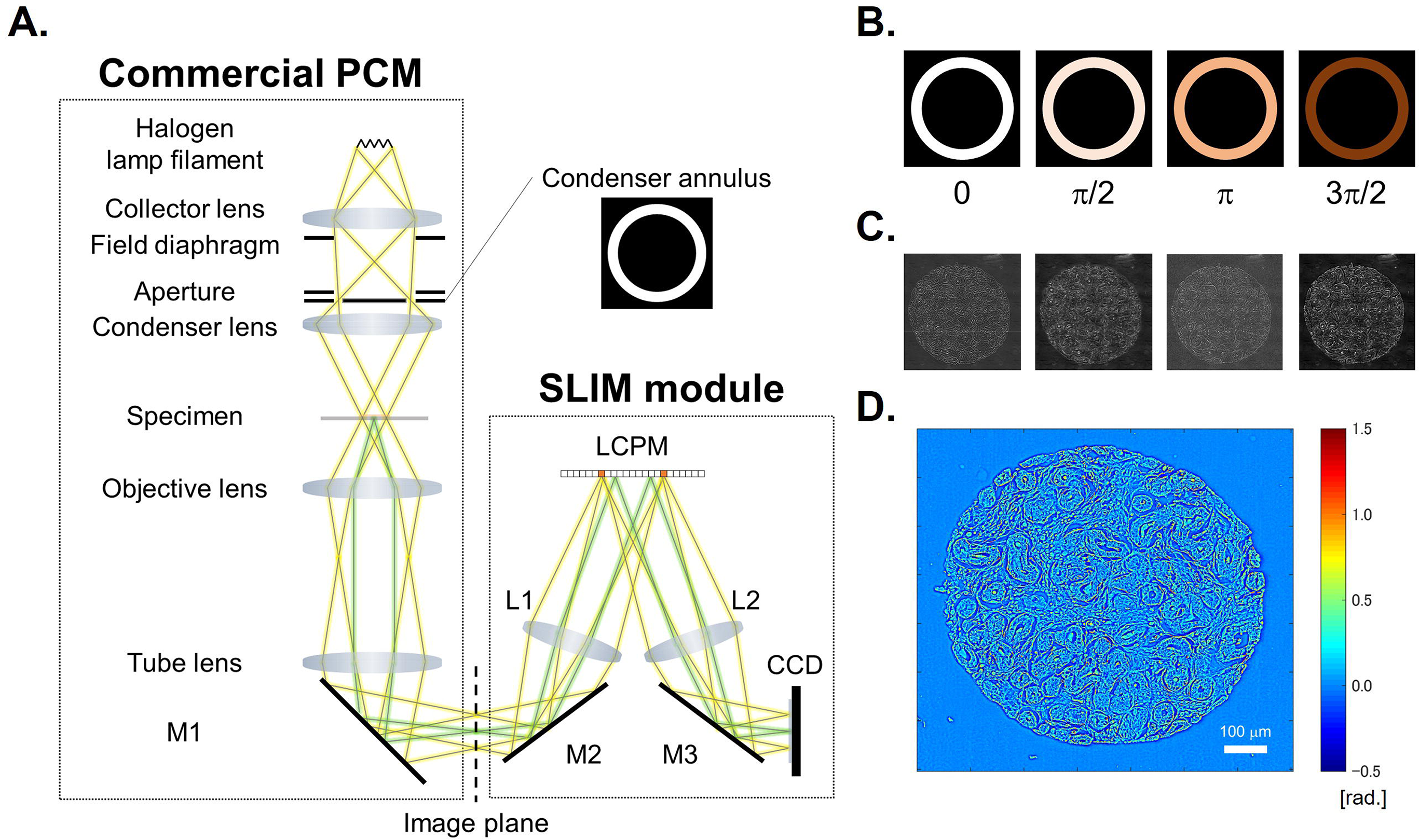
SLIM system. (A) Schematic setup. (B) The phase rings and (C) their corresponding intensity images captured by CCD. (D) The retrieved quantitative phase image.

**Fig 2.**
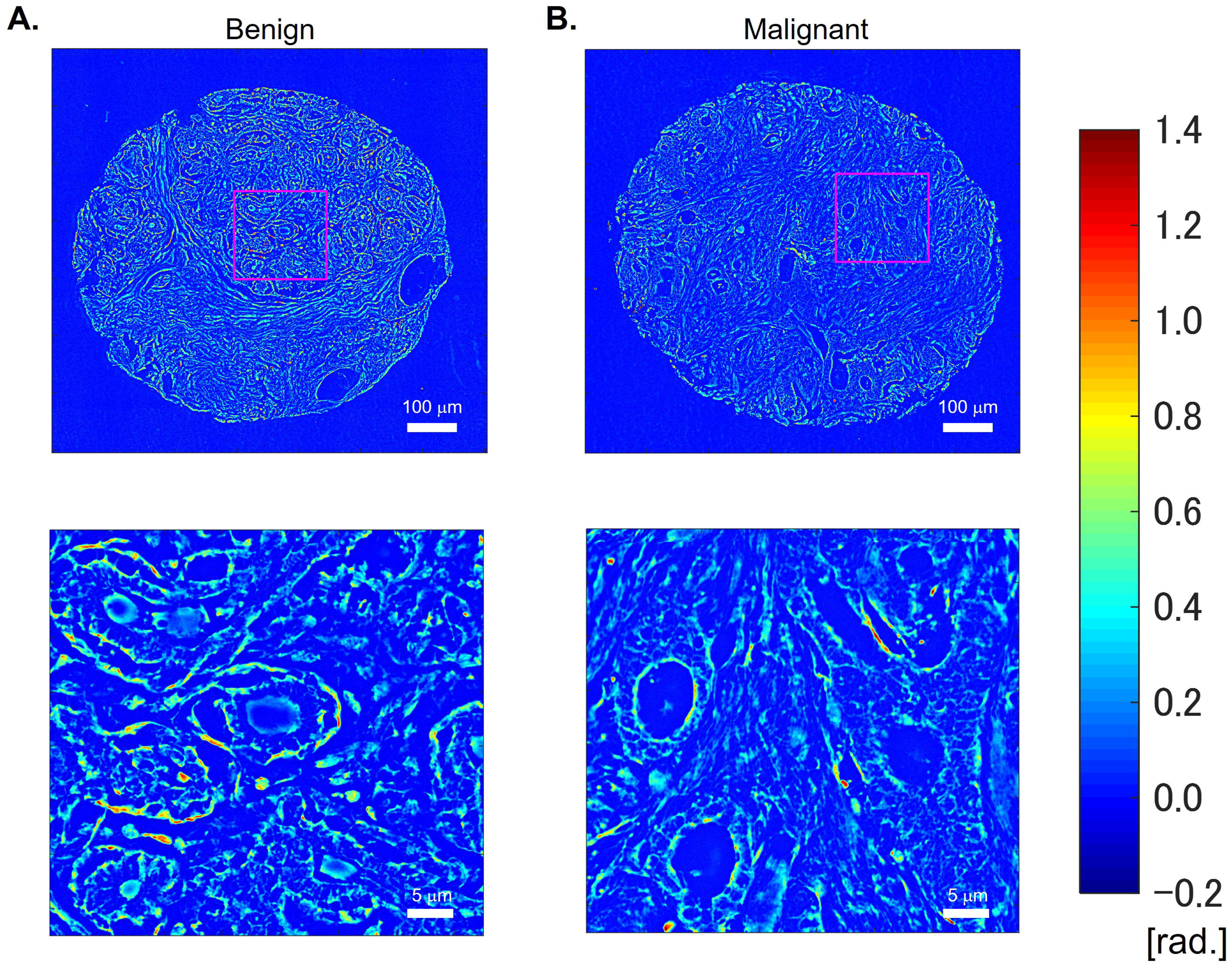
Quantitative phase images and their enlarged view. (A) Benign and (B) malignant breast tissues.

By definition, the disorder strength map, *L_d_*(*x*, *y*), is expressed as:

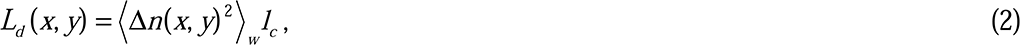

Here, *<…>_w_* denotes the average within the window of interest, Δ means the difference from its average, i.e., Δ*n*(*x*, *y*) = *n*(*x*, *y*) - *<n*(*x*, *y*)*>_w_*, and *l_c_* is the spatial autocorrelation length. Figure 3A shows the quantitative phase image (*x*, *y*), which contains information about the spatial variation of the refractive index change of tissues as expressed by Eq. 1. The local variance and average of the phase has the form, respectively,

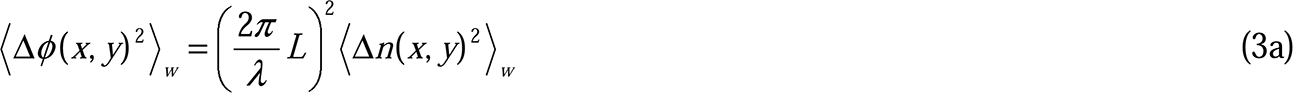
and

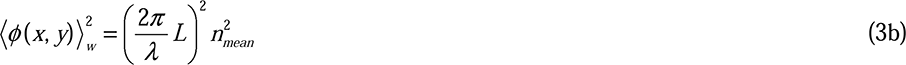

**Fig 3.**
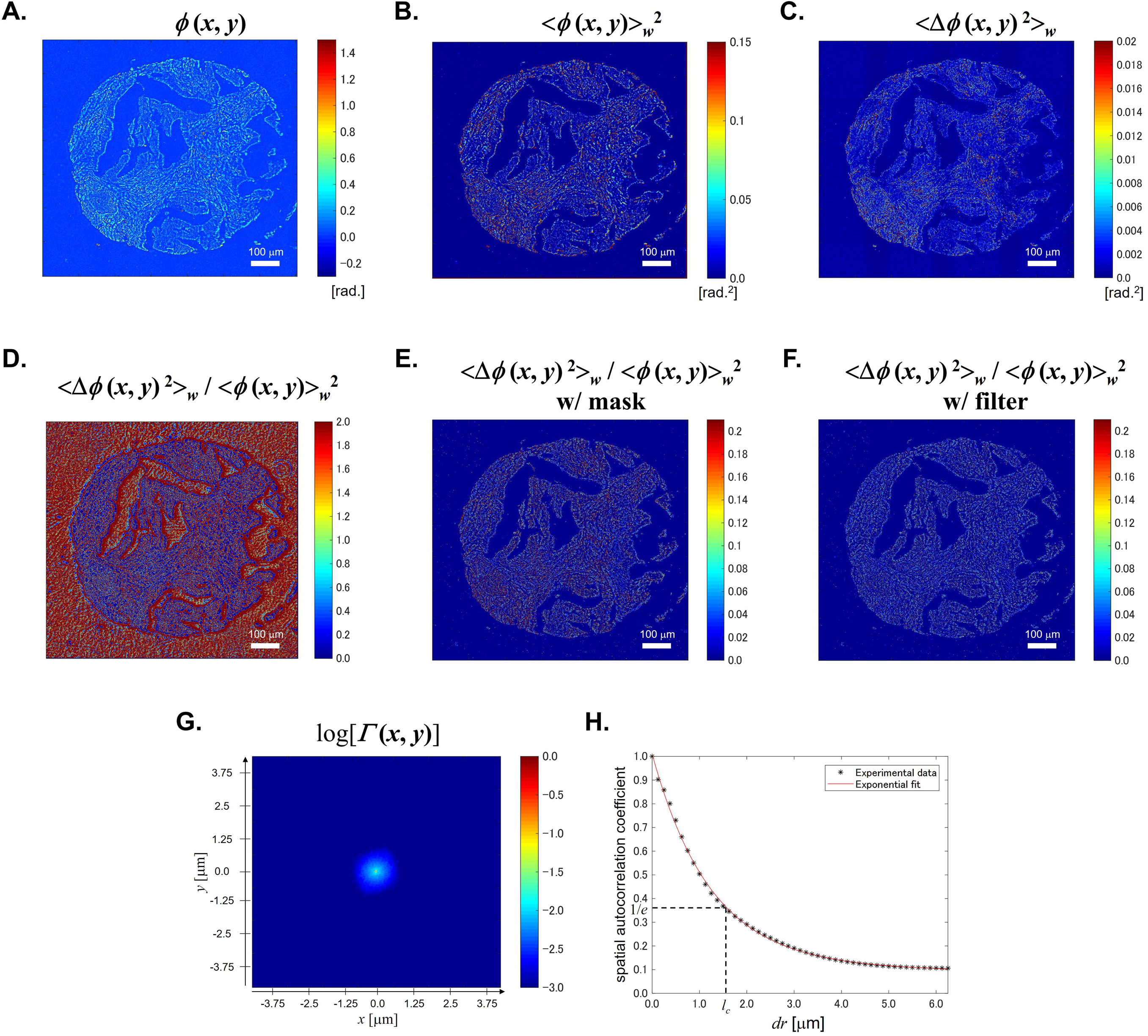
Procedures. (A) Original phase image. (B) Local phase average map. (C) Local phase variance map. (D) Local phase fluctuation map. (E) Local phase fluctuation map with background reduction mask. (F) Local phase fluctuation map with edge reduction filter. (G) 2D spatial autocorrelation. (H) 1D spatial autocorrelation.

Here, *n_mean_* is the average of the refractive index in the tissue. Thus, the local refractive index fluctuation map, which is independent of the thickness, can be computed as

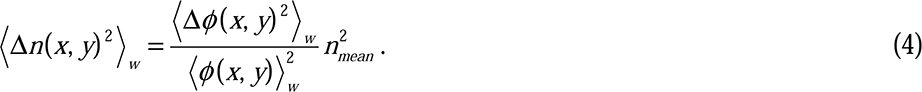

Therefore, we can rewrite Eq. 2 and obtain the final form to calculate the disorder strength map from the quantitative phase image as

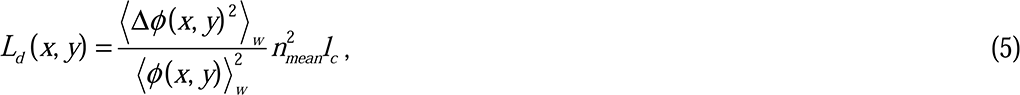

In our calculation, we used a window of 5×5 pixels (0.125 μm/pixel). Figure 3B and 3C show the example of the calculation result of < (*x*, *y*)*>_w_*^2^ and <Δ (*x*, *y*)^2^*>_w_*, respectively. Also, from these two images, we can obtain <Δ (*x y*)^2^*>_w_* / < (*x*, *y*)*>_w_^2^* as shown in Fig. 3D. Since our interest is the fluctuation only in the tissue region, the background pixels were excluded. In our calculation, the pixels which satisfy < (*x*, *y*)*>_w_* < 0.075 rad. are treated as background pixels. Here, we note that the selected threshold value of 0.075 rad. is about 3 times larger than the standard deviation of (*x*, *y*) in an arbitrary selected background region. Figure 3E shows <Δ (*x*, *y*)^2^*>_w_* / < (x *y*)*>_w_*^2^ after applying this mask. Furthermore, because the very large value of <Δ (*x*, *y*)^2^*>_w_*/ ***<*** (*x*, *y*)*>_w_^2^* are observed in the small area including the edge of the tissue where the thickness in the area might not be constant, the pixels having the value larger than 0.21 are filtered out as shown in Fig. 3F. Consequently, the disorder strength can be calculated using the resulting phase image. Figure 3G shows the normalized 2D spatial autocorrelation function of (*x*, *y*), *Γ*(*x*, *y*). From this, we re-plot the normalized spatial autocorrelation function in terms of *r* = (*x*^2^ + *y*^2^)^1/2^ as shown in Fig. 3H. Then, the spatial autocorrelation length, *l_c_*, is defined as the half width at 1/*e* of the maximum, i.e., *Γ(r* = *l_c_)* = 1/*e*.

## Results

Figures 4A and 4B shows the disorder strength maps of benign and malignant samples, respectively. It can be seen that the disorder strength in the malignant sample is larger than that in the benign sample. Also, we can find that the disorder strength obtained in our calculation is of the order of 1 μm which is higher than that obtained in previous studies [21, 32]. This is likely caused by the difference in contrast between different techniques. Laser QPI methods are known to produce lower contrast phase images, (*x*, *y*). As a result, Δ (*x*, *y*) has much smaller values in this case. Furthermore, using smaller windows for averaging also results in lower Δ (*x*, *y*) values. Images obtained by SLIM has higher contrast than those obtained by other imaging systems, which result in higher phase variance. Figure 5 shows the averaged disorder strength across the tissue area. The error bar in the figure denotes the standard error in 20 samples each. The p-value between the benign and malignant samples using two-sided Wilcoxon ranksum test was 0.0066. The results indicate that there are statistically significant differences between these two groups, therefore, the disorder strength can be utilized as a marker for a tissue screening. As demonstrated in [32], measurement of disorder strength can inform on the mechanical properties of a biological specimen and our results motivate further studies investigating how tissue mechanics vary with disease progression, with disorder strength used as a convenient metric.

**Fig 4.**
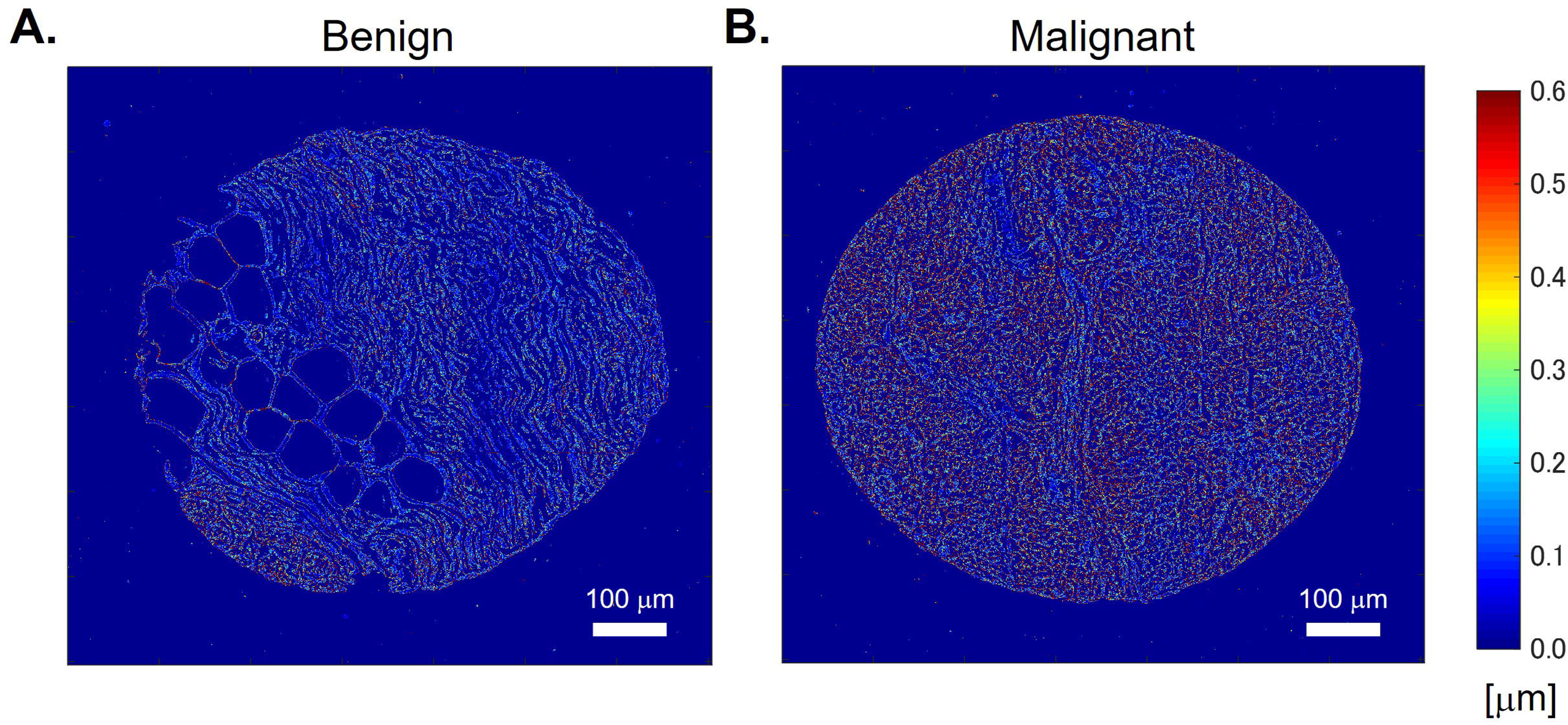
Disorder strength maps. (A) Benign and (B) malignant tissues.

**Fig 5.**
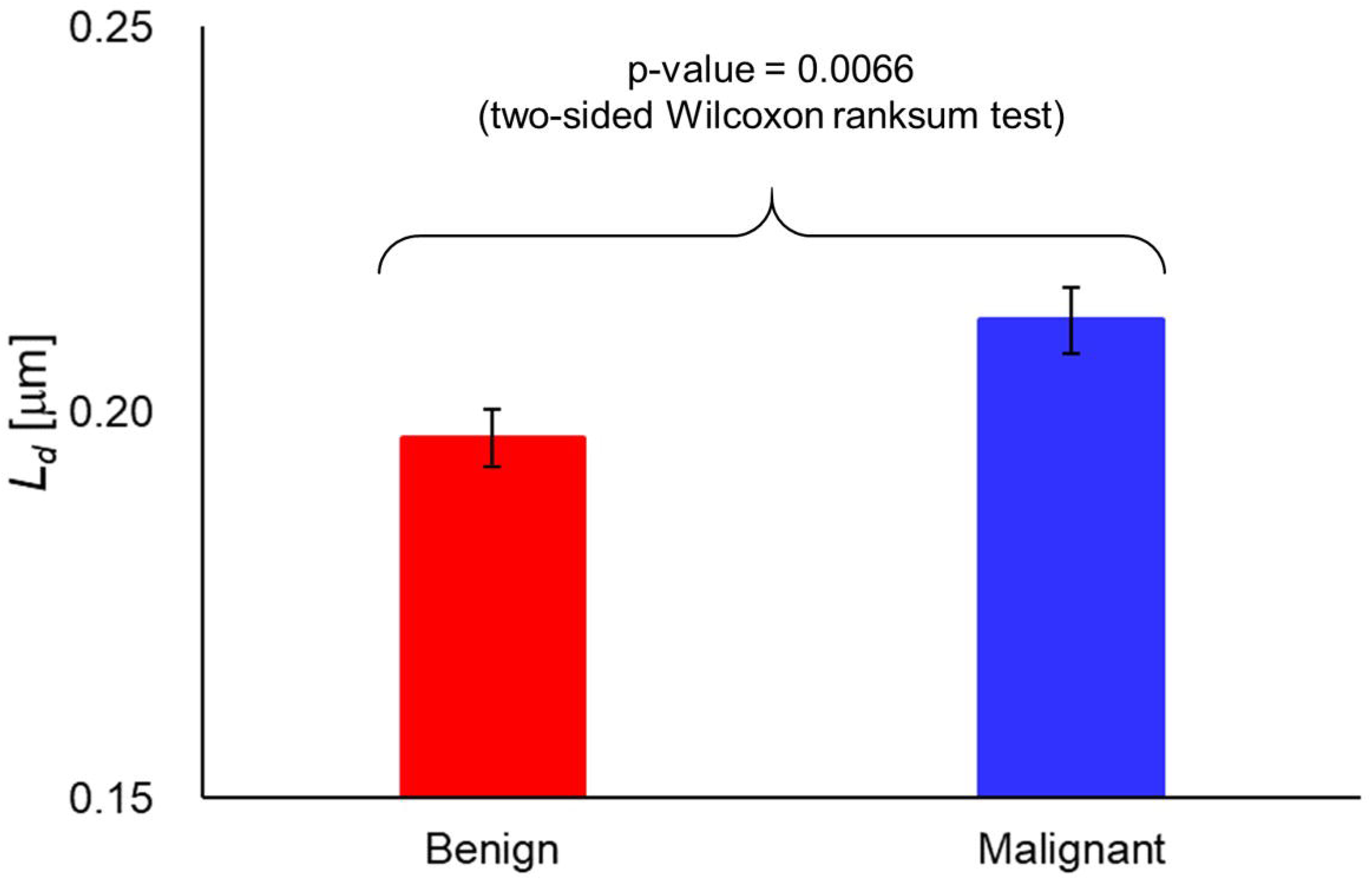
Disorder strength of benign (N=20) and malignant (N=20) tissues.

## Summary

In summary, we showed that the disorder strength measured from QPI is a quantitative marker of malignancy that can be used to classify benign and malignant breast cores. This marker, which measures the tissue “disorder strength”, via the refractive index fluctuations is obtainable from unlabeled tissue images, meaning that it is not affected by stain variation among different samples.

Finally, we note that previous publications have shown that, from quantitative phase images of tissue slices, one can extract the scattering mean free path and anisotropy factor of the bulk [33, 34]. This result, known as the scattering phase theorem, has led to multiple SLIM studies of using scattering parameters for diagnosis and prognosis [10, 12–15, 35]. Specifically, the scattering mean free path, *l_s_* relates to the phase variance as *l_s_* = *L* / <Δ (x *y*)^2^*>_w_* [34]. Therefore, from Eq. 5, we see that the disorder strength and scattering mean free path are simply inversely proportional, *L_d_ ∝* 1 / *l_s_*. The physical significance of this result is straight forward: higher tissue disorder generates stronger scattering, which implies shorter *l_s_*. It is, thus, not surprising that both parameters have been used successfully in cancer pathology, as they both report on tissue inhomogeneity. SLIM provides a robust, high-throughput approach to imaging histology slides. Recent advances in SLIM data acquisition allowed us to image an entire microscope slide, containing hundreds of tissue cores, at 0.5 μm transverse resolution in 45 min, while maintaining the sub-nanometer path-length sensitivity [16]. Because SLIM can be implemented as an upgrade of the existing microscopes, we anticipate that it can be plugged into the existing pathology work flow and help solve many problems of clinical importance.

## Acknowledgements

This work was supported by the National Science Foundation (CBET-0939511 STC, DBI 14–50962 EAGER, IIP-1353368, CBET-1040461 MRI). G. P. has financial interest in Phi Optics, Inc., a company that develops quantitative phase imaging technologies. For more information, visit http://light.ece.illinois.edu/.

We would like to thank Prof. Garth H. Rauscher for pathology expertise.

## References

1. Sanduleanu S, Driessen A, Gomez-Garcia E, Hameeteman W, de Bruïne A, Masclee A. In vivo diagnosis and classification of colorectal neoplasia by chromoendoscopy-guided confocal laser endomicroscopy.

2. Cancer Fact Sheets: Breast Cancer: International Agency for Research on Cancer -World Health Organization. Available from: http://gco.iarc.fr/today/fact-sheets-cancers?cancer=15&type=0&sex=2.

3. Kumar GL, Kiernan JA, Dako AS. Education Guide -Special Stains and H & E: Pathology: Dako North America; 2010.

4. Popescu G. Quantitative Phase Imaging of Cells and Tissues: McGraw Hill; 2011.

5. Wang Z, Millet L, Mir M, Ding H, Unarunotai S, Rogers J, et al. Spatial light interference microscopy (SLIM). Optics Express. 2011;19:1016–26.

6. Wang P, Bista R, Rohit B, Brand E, Randall, Liu Y. Spatial-domain low-coherence quantitative phase microscopy for cancer diagnosis Optics Letters. 2010 35(17):2840. Epub 2842.

7. Uttam S, Pham HV, LaFace J, Leibowitz B, Yu J, Brand RE, et al. Early Prediction of Cancer Progression by Depth-Resolved Nanoscale Mapping of Nuclear Architecture from Unstained Tissue Specimens. Cancer Research. 2015;75(22):4718–27. doi: 10.1158/0008–5472.can-15-1274.

8. Mir M, Wang Z, Shen Z, Bednarz M, Bashir R, Golding I, et al. Optical measurement of cycle-dependent cell growth,. Proceedings of the National Academy of Sciences. 2011;108.

9. Cooper KL, Oh S, Sung Y, Dasari RR, Kirschner MW, Tabin CJ. Multiple phases of chondrocyte enlargement underlie differences in skeletal proportions. Nature. 2013;495(7441):375–8. Epub 2013/03/15. doi: 10.1038/nature11940. PubMed PMID: 23485973; PubMed Central PMCID: PMCPMC3606657.

10. Wang Z, Tangella K, Balla A, Popescu G. Tissue refractive index as marker of disease. J Biomed Opt. 2011;16(11):116017. doi: 10.1117/1.3656732. PubMed PMID: 22112122; PubMed Central PMCID: PMC3223513.

11. Park Y, Diez-Silva M, Popescu G, Lykotrafitis G, Choi W, Feld MS, et al. Refractive index maps and membrane dynamics of human red blood cells parasitized by Plasmodium falciparum. Proceedings of the National Academy of Sciences. 2008;105.

12. Sridharan S, Macias V, Tangella K, Kajdacsy-Balla A, Popescu G. Prediction of Prostate Cancer Recurrence Using Quantitative Phase Imaging. Scientific Reports. 2015;5:9976. doi: 10.1038/srep09976 http://www.nature.com/articles/srep09976#supplementary-information.

13. Majeed H, Kandel ME, Han K, Luo Z, Macias V, Tangella K, et al. Breast cancer diagnosis using spatial light interference microscopy. Journal of Biomedical Optics. 2015;20(11):111210-. doi: 10.1117/1.JBO.20.11.111210.

14. Majeed H, Sridharan S, Mir M, Ma L, Min E, Jung W, et al. Quantitative phase imaging for medical diagnosis. J Biophotonics. 2017;10(2):177–205. Epub 2016/08/20. doi: 10.1002/jbio.201600113. PubMed PMID: 27539534.

15. Nguyen TH, Sridharan S, Macias V, Kajdacsy-Balla A, Melamed J, Do MN, et al. Automatic Gleason grading of prostate cancer using quantitative phase imaging and machine learning. J Biomed Opt. 2017;22(3):36015. Epub 2017/03/31. doi: 10.1117/1.JBO.22.3.036015. PubMed PMID: 28358941.

16. Kandel ME, Sridharan S, Liang J, Luo Z, Han K, Macias V, et al. Label-free tissue scanner for colorectal cancer screening. Journal of Biomedical Optics. 2017;22(6):066016-. doi: 10.1117/1.JBO.22.6.066016.

17. Bista RK, Wang P, Bhargava R, Uttam S, Hartman DJ, Brand RE, et al. Nuclear nano-morphology markers of histologically normal cells detect the "field effect" of breast cancer. Breast Cancer Res Treat. 2012;135(1):115–24. Epub 2012/06/19. doi: 10.1007/s10549–012-2125-2. PubMed PMID: 22706633; PubMed Central PMCID: PMCPMC3566261.

18. Wang P, Bista RK, Khalbuss WE, Qiu W, Uttam S, Staton K, et al. Nanoscale nuclear architecture for cancer diagnosis beyond pathology via spatial-domain low-coherence quantitative phase microscopy.

19. Liu Y, Uttam S, Alexandrov S, Bista RK. Investigation of nanoscale structural alterations of cell nucleus as an early sign of cancer. BMC Biophysics. 2014;7(1):1. doi: 10.1186/2046–1682-7-1.

20. Pham HV, Pantanowitz L, Liu Y. Quantitative phase imaging to improve the diagnostic accuracy of urine cytology. Cancer cytopathology. 2016;124(9). Epub 650.

21. Subramanian H, Pradhan P, Liu Y, Capoglu IR, Rogers JD, Roy HK, et al. Partial-wave microscopic spectroscopy detects subwavelength refractive index fluctuations: an application to cancer diagnosis. Optics Letters. 2009;34.

22. Subramanian H, Roy HK, Pradhan P, Goldberg MJ, Muldoon J, Brand RE, et al. Nanoscale Cellular Changes in Field Carcinogenesis Detected by Partial Wave Spectroscopy. Cancer research. 2009;69(13):5357–63. doi: 10.1158/0008–5472.CAN-08-3895. PubMed PMID: PMC2802178.

23. Roy HK, Subramanian H, Damania D, Hensing TA, Rom WN, Pass HI, et al. Optical Detection of Buccal Epithelial Nanoarchitectural Alterations in Patients Harboring Lung Cancer: Implications for Screening. Cancer research. 2010;70(20):7748–54. doi: 10.1158/0008–5472.CAN-10-1686. PubMed PMID: PMC3703950.

24. Damania D, Subramanian H, Tiwari AK, Stypula Y, Kunte D, Pradhan P, et al. Role of Cytoskeleton in Controlling the Disorder Strength of Cellular Nanoscale Architecture. Biophysical Journal. 2010;99(3):989–96. doi: 10.1016/j.bpj.2010.05.023. PubMed PMID: PMC2913198.

25. Jun Soo Kim and Prabhakar Pradhan and Vadim Backman and Igal S. The influence of chromosome density variations on the increase in nuclear disorder strength in carcinogenesis. Physical Biology. 2011;8(1):015004.

26. Damania D, Roy HK, Subramanian H, Weinberg DS, Rex DK, Goldberg MJ, et al. Nanocytology of rectal colonocytes to assess risk of colon cancer based on field cancerization. Cancer research. 2012;72(11):2720–7. doi: 10.1158/0008–5472.CAN-11-3807. PubMed PMID: PMC3557939.

27. Stypula-Cyrus Y, Damania D, Kunte DP, Cruz MD, Subramanian H, Roy HK, et al. HDAC Up-Regulation in Early Colon Field Carcinogenesis Is Involved in Cell Tumorigenicity through Regulation of Chromatin Structure. PLoS ONE. 2013;8(5):e64600. doi: 10.1371/journal.pone.0064600. PubMed PMID: PMC3665824.

28. Chandler JE, Subramanian H, Maneval CD, White CA, Levenson RM, Backman V. High-speed spectral nanocytology for early cancer screening. Journal of Biomedical Optics. 2013;18(11):117002. doi: 10.1117/1.JBO.18.11.117002. PubMed PMID: PMC3817856.

29. Konda VJA, Cherkezyan L, Subramanian H, Wroblewski K, Damania D, Becker V, et al. Nanoscale markers of esophageal field carcinogenesis: potential implications for esophageal cancer screening. Endoscopy. 2013;45(12):983–8. doi: 10.1055/s-0033–1344617. PubMed PMID: PMC4195538.

30. Roy HK, Damania DP, DelaCruz M, Kunte DP, Subramanian H, Crawford SE, et al. Nano-architectural Alterations in Mucus Layer Fecal Colonocytes in Field Carcinogenesis: Potential for Screening. Cancer prevention research (Philadelphia, Pa). 2013;6(10):10.1158/940–6207.CAPR-13-0138. doi: 10.1158/1940-6207.CAPR-13-0138. PubMed PMID: PMC3873747.

31. Roy HK, Brendler CB, Subramanian H, Zhang D, Maneval C, Chandler J, et al. Nanocytological Field Carcinogenesis Detection to Mitigate Overdiagnosis of Prostate Cancer: A Proof of Concept Study. PLOS ONE. 2015;10(2):e0115999. doi: 10.1371/journal.pone.0115999.

32. Eldridge WJ, Steelman ZA, Loomis B, Wax A. Optical Phase Measurements of Disorder Strength Link Microstructure to Cell Stiffness. Biophysical Journal. 2017;112(4):692–702. doi: https://doi.org/10.1016/j.bpj.2016.12.016.

33. Ding H, Wang Z, Liang X, Boppart S, Tangella K, Popescu G. Measuring the scattering parameters of tissues from quantitative phase imaging of thin slices. Optics letters. 2011;36(12):2281.

34. Wang Z, Ding H, Popescu G. Scattering-phase theorem. Optics letters. 2011;36(7):1215.

35. Majeed H, Okoro C, Kajdacsy Balla A, Toussaint K, Popescu G. Quantifying collagen fiber orientation in breast cancer using quantitative phase imaging. Journal of biomedical optics. 2017;22(4):046004.

